# Gestational age at birth and risk of intellectual disability without a common genetic cause: findings from the Stockholm Youth Cohort

**DOI:** 10.1101/129049

**Authors:** Hein Heuvelman, Kathryn Abel, Susanne Wicks, Renee Gardner, Edward Johnstone, Brian Lee, Cecilia Magnusson, Christina Dalman, Dheeraj Rai

**Author notes:** **Correspondence to:** Hein Heuvelman. Centre for Academic Mental Health, School of Social and Community Medicine, University of Bristol, Oakfield House, Oakfield Grove, BS8 2BN, Bristol, UK. **Email addresses of other authors:** Kathryn Abel; Susanne Wicks; Renee Gardner; Edward Johnstone; Brian Lee; Cecilia Magnusson; Christina Dalman; Dheeraj Rai. **Ethical approval:** Ethical approval for this study was granted by the research ethics committee at Karolinska Institute [2010/1185-31/5 and 2013/1118-32], allowing record linkage without personal consent when the confidentiality of the individuals is maintained. The personal identity of participants was replaced with a serial number before the research group were given access to these data. It is of paramount importance to ensure the protection of the personal integrity against any violations, and legislation regulating the handling of information that is directly or indirectly linked to a person is in place (the Personal Data Act).

## Abstract

**Background:** Preterm birth is linked to intellectual disability and there is evidence to suggest post-term birth may also incur risk. However, these associations have not yet been investigated in the absence of common genetic causes of intellectual disability (where risk associated with late delivery may be preventable) or with methods allowing stronger causal inference from non-experimental data. We aimed to examine risk of intellectual disability without a common genetic cause across the entire range of gestation, using a matched-sibling design to account for unmeasured confounding by shared familial factors.

**Methods and Findings:** We conducted a population-based retrospective study using data from the Stockholm Youth Cohort (n=499,621) and examined associations in a nested cohort of matched siblings (n=8,034). Children born at non-optimal gestational duration (before/after 40 weeks 3 days) were at greater risk of intellectual disability. Risk was greatest among those born extremely early (adjusted OR_24 weeks_=14.54 [95% CI 11.46–18.44]), lessening with advancing gestational age toward term (aOR_32 weeks_=3.59 [3.22–4.01]; aOR_37 weeks_=1.50 [1.38–1.63]); aOR_38 weeks_=1.26 [1.16-1.37]; aOR_39_ weeks=1.10 [1.04-1.17]) and increasing with advancing gestational age post-term (aOR_42 weeks_=1.16 [1.08–1.25]; aOR_42 weeks_=1.41 [1.21–1.64]; aOR_44 weeks_=1.71 [1.34–2.18]; aOR_45 weeks_=2.07 [1.47–2.92]). Associations persisted in a nested cohort of matched outcome-discordant siblings suggesting they were robust against confounding from shared genetic or environmental traits, although there may have been residual confounding by unobserved non-shared characteristics. Risk of intellectual disability was greatest among children showing evidence of fetal growth restriction, especially when birth occurred before or after term.

**Conclusions:** Birth at non-optimal gestational duration may be linked causally with greater risk of intellectual disability. The mechanisms underlying these associations need to be elucidated as they will be relevant to clinical practice concerning elective delivery within the term period and the mitigation of risk in children who are born post-term.

## Introduction

Intellectual disability is a group of developmental disorders evident early in childhood and characterized by cognitive and functional impairments as a result of delayed or incomplete development of the mind^[1]^. Individuals with intellectual disability have a reduced ability to understand new or complex information and to learn and apply new skills, resulting in a reduced ability to cope independently^[2]^. Intellectual disability is thought to affect over 1 percent of the population^[3, 4]^ although estimates vary with the demographic and socioeconomic composition of study populations^[4, 5]^ and with definitions and study design^[5, 6]^. The cost of intellectual disability to individuals and society is substantial^[7]^ and people living with these disabilities often face significant stigma^[8]^ while encountering substantial health and social inequalities and early mortality^[9]^.

Although there are many risk factors, a specific cause is identified for less than half of those with mild disabilities (IQ range 50-69) who make up the majority of cases^[3, 10]^. Mild intellectual disability often clusters within families^[10]^ suggesting that genetic or other shared familial factors may influence risk. When disabilities are more severe, specific causes are identified in over 75 percent of cases, often involving genetic or chromosomal abnormalities and inborn errors of metabolism [10]. When intellectual disability is present without a specific genetic or chromosomal cause it is associated with advanced maternal age, maternal risk behaviors or medical problems during pregnancy and fetal growth restriction^[11]^, suggesting that these may be risk factors.

While it is known that children born preterm (<37 completed weeks) are at greater risk of intellectual disability than those born at term^[12]^, less is known about the development of risk along the gestational course, or about risk among post-term children (>41 weeks). This is important considering there is increasing evidence to suggest that post-term birth is associated with cognitive and academic deficits in childhood and adolescence^[13-16]^, especially when the baby is growth-restricted^[16]^.

The association between the full range of gestational duration, from very early to very late births, and intellectual disability has not yet been examined in population-based studies. Furthermore, the evidence to date is insufficient because of incomplete control of confounding from shared familial factors and insufficient recognition that genetic causes of intellectual disability may also influence gestational duration^[17, 18]^.

Therefore, in a large Swedish population-based cohort, we aimed to: 1) examine the associations between gestational age and intellectual disability without a common genetic cause, taking into account a range of potential confounders; 2) examine interactions between gestational duration and fetal growth in relation to risk of intellectual disability; and 3) explore the causal nature of associations between gestational duration and risk of intellectual disability in a nested cohort of matched outcome-discordant siblings.

## Methods

### Study cohort

The Stockholm Youth cohort is a register-based cohort of all individuals who lived in Stockholm County for at least one year between 2001 and 2011 and were aged between 0 and 17 years during that period (n=736,180)^[19]^. Using unique personal identification numbers, cohort members and their first degree relatives were linked with a range of national and regional registers including information on pregnancy- and birth related characteristics, socioeconomic characteristics and medical and psychiatric diagnoses.

We excluded individuals with genetic and inborn metabolic syndromes who had been diagnosed with intellectual disability (13.6 % of cases in our study population), children born outside Sweden, multiple births, adoptees, children <4 years of age by the end of follow up on the 31^st^ of December 2011, with a missing link to biological parents, or with missing data on gestational age or other covariates (see Figure 1). We excluded individuals with implausible combinations of gestational age and birth weight following methods described elsewhere^[16]^. This left a cohort of 499,621 individuals to examine population-level associations between gestational age and intellectual disability. To examine associations among matched siblings, we excluded individuals without full siblings in the cohort and families with outcome-concordant offspring (n=491,587) leaving a cohort of 8,034 matched outcome-discordant full siblings.

**Fig 1:**
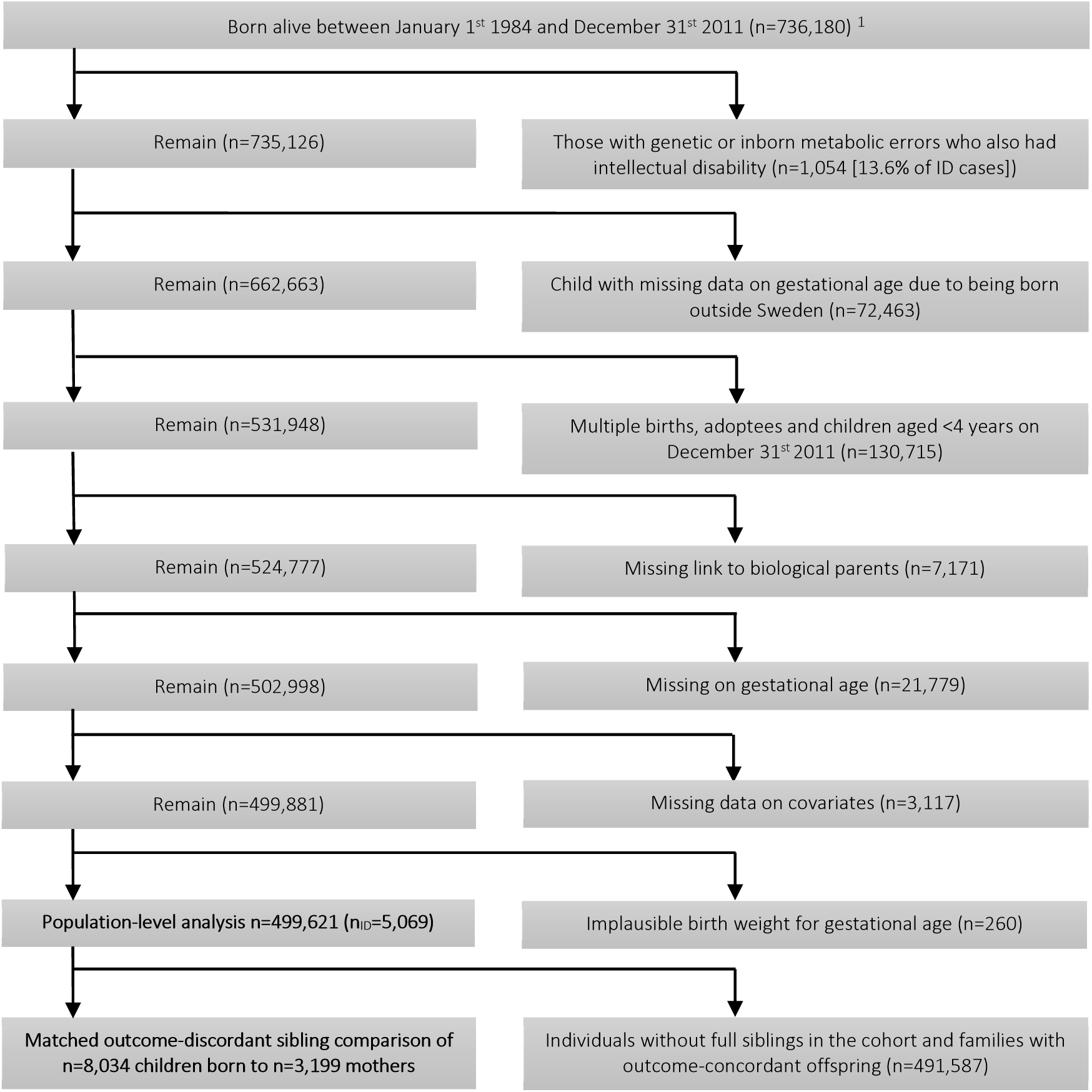
Selection of the study cohort.

### Exposure

We obtained information on gestational age at birth from the Medical Birth Register (MBR), constructing a categorical variable to define extremely to very preterm births (21-31 completed weeks), moderately to late preterm births (32-36 weeks), term births (37-41 weeks), post-term births (42 weeks) and very post-term births (43-45 weeks) for use in descriptive statistics and as an exposure variable in regression analyses. We also used a continuous definition of gestational age (in days) for regression analyses. A measure of weight-for-gestational age was calculated using week- and sex-specific birth weight distributions, identifying individuals in the lower and upper deciles as born small or large for gestational age respectively. To examine interactions between gestational duration and fetal growth, we constructed a categorical variable to identify those born preterm (<37 weeks) and small for gestational age, appropriate for gestational age (11^th^ centile to 90^th^ centile) or large for gestational age; those born at term (37-41 weeks) and small, appropriate or large for gestational age; and those born post-term (≥42) and small, appropriate or large for gestational age.

### Outcome

We used a multisource ascertainment approach to identify cohort members with intellectual disability, similar to the case identification for autism described elsewhere ^[19]^. We used the national patient register, the Stockholm county child and adolescent mental health register, the Stockholm country healthcare database (VAL) and the Stockholm adult psychiatry register to identify inpatient or outpatient diagnoses of intellectual disability recorded using ICD-10 (F70-79) and DSM-IV (317-318) codes and supplemented these diagnoses with a record of care at specialist habilitation services for individuals with intellectual disability in Stockholm County. We identified individuals with genetic defects and inborn errors of metabolism commonly associated with intellectual disability to identify cases where a known genetic or metabolic cause was present (Table S1).

### Covariates

To control for secular change in obstetric and diagnostic practice, we obtained year of birth from the Medical Birth Register (MBR). We then identified additional covariates which in the literature have been associated with pregnancy duration and risk of intellectual disability in offspring. From the MBR, we extracted data for offspring sex^[5, 20]^, parity (1/ 2/ 3/ 4+)^[11, 21]^, birth weight^[11]^, maternal age (<20/ 20-24/ 25-29/ 30-34/ 35-39/ 40-44/ 45+)^[11, 21]^, gestational diabetes^[11, 22]^ and gestational hypertension or preeclampsia^[11, 23]^. We also extracted information for maternal and paternal country of birth (Sweden/ other Nordic/ other European/ Russia or Baltic States / Africa /Middle East/ Asia or Oceania/ North America/ South America)^[5, 24]^, maternal and paternal history of psychiatric treatment^[25, 26]^, quintiles of disposable family income adjusted for inflation and family size^[27, 28]^, and parental educational attainment (≤9 years/ 10-12 years/ ≥13 years)^[28, 29]^ at (or as close as possible to) birth.

### Statistical analyses

Analyses were conducted in Stata/MP version 14.2. We examined the characteristics of the study cohort by gestational duration at birth. To examine population-level associations between gestational duration and risk of intellectual disability, we used generalized estimating equation (GEE) multivariable regression models with a logit link function, exchangeable correlation structure and robust variance estimators to ensure that the standard errors of our estimates were robust against clustering of intellectual disability within families^[30]^. We calculated restricted cubic regression splines based on five knot locations (5^th^, 27^th^, 50^th^, 73^rd^ and 95^th^ percentiles of the gestational age distribution) to allow for non-linear associations between continuously varying gestational duration and later risk of intellectual disability^[31]^. We statistically adjusted our estimates for covariates and calculated odds ratios by continuously varying gestational age at birth to estimate risk of intellectual disability associated with birth at specific moments along the gestational course. We investigated potential interactions between gestational age and fetal growth using GEE models with a categorical exposure variable to assess risk of intellectual disability among those born at varying gestational duration (preterm/ term/ post-term) and weight for gestational age (small/ appropriate/ large) with statistical adjustment for confounders.

In a nested cohort of matched outcome-discordant siblings we examined associations between continuously varying gestational age and risk of intellectual disability with conditional likelihood logistic regression models. This allowed us to explore the potential influence of unobserved familial traits, e.g. residual genetic risk/ unmeasured socioeconomic factors/ parental health behaviors, which may have confounded associations between gestational length and risk of intellectual disability. If we were to observe associations at the population level, non-association within families would suggest confounding by these shared familial traits. Conversely, replication of population-level associations within families would suggest they were robust against shared familial confounding, thereby allowing stronger causal inference from our result ^[32]^. We statistically adjusted within-family associations for non-shared confounding characteristics including sex, parity, gestational diabetes, hypertension or preeclampsia, weight for gestational age, maternal and paternal age, disposable family income quintile, and parental educational attainment.

### Sensitivity analyses

We compared characteristics for those with missing and complete data to assess whether our estimates may have been affected by selection bias (Table S2). To ensure that the association between gestational age and intellectual disability was not driven by presence of co-occurring autism spectrum disorder or attention deficit hyperactivity disorder (which are associated with intellectual disability ^[33-35]^ and for which risk may also vary by gestational age ^[36, 37]^) we examined associations in a subset of the cohort without a record of these conditions (Figure S1 and Table S3). We examined whether the risks of intellectual disability associated with preterm or post-term birth varied with mode of delivery (Tables S4 and S5) using categorical measures to identify those born vaginally or by Caesarean section and in unassisted or forceps-/ ventouse-assisted deliveries at varying gestational duration. Finally, we conducted post-hoc analyses to assess whether risk varied among children born in spontaneous or induced deliveries at varying gestational duration (Table S6).

## Results

Prevalence of intellectual disability without a common genetic cause was estimated at 1% in our study population (Figure 1). Characteristics of the study cohort are described in Table 1. Prevalence among those born at term gestation was 0.9 percent. By contrast, 5.6 percent of children born extremely to very preterm and 1.6 percent of those born very post-term had intellectual disability.

**Table 1:** Characteristics of the sample by exposure status.

|  |  | Extremely<br>to very<br>preterm | Moderate<br>to late<br>preterm | Term | Post-term | Very<br>Post-term |
| --- | --- | --- | --- | --- | --- | --- |
| Gestational weeks |  | 21 - 31 | 32 - 36 | 37 - 41 | 42 | 43 - 45 |
| Number of observations |  | 2,601 | 20,271 | 438,215 | 34,828 | 3,706 |
| Percentage of the cohort |  | 0.5 | 4.1 | 87.7 | 7.0 | 0.7 |
|  |  | % | % | % | % | % |
| Female child |  | 45.2 | 46.3 | 49.3 | 44.1 | 43.9 |
| Mother's number of prior pregnancies | 0 | 53.4 | 53.4 | 44.2 | 53.8 | 61.5 |
|  | 1 | 27.1 | 28.4 | 37.0 | 30.1 | 23.5 |
|  | 2 | 12.5 | 11.8 | 13.6 | 11.6 | 10.0 |
|  | 3+ | 7.0 | 6.4 | 5.3 | 4.5 | 5.0 |
| Birth weight in grams | <2500 | 99.7 | 37.6 | 1.1 | 0.1 | 0.1 |
|  | 2500 – 4500 | 0.4 | 62.5 | 98.6 | 98.5 | 98.0 |
|  | >4500 | 0.0 | 0.0 | 0.3 | 1.4 | 1.9 |
| Gestational hypertension or preeclampsia |  | 23.6 | 12.9 | 3.2 | 1.9 | 1.5 |
| Gestational diabetes |  | 1.5 | 1.7 | 0.8 | 0.4 | 0.4 |
| Delivery by Caesarean Section |  | 58.9 | 32.4 | 13.3 | 16.5 | 22.5 |
| Delivery assisted with ventouse or forceps |  | 1.4 | 4.7 | 8.0 | 14.1 | 14.9 |
| Maternal psychiatric history |  | 38.6 | 37.0 | 32.7 | 31.5 | 33.0 |
| Paternal psychiatric history |  | 23.7 | 22.6 | 20.9 | 20.4 | 21.2 |
| Mother's country of birth | Sweden | 72.2 | 75.2 | 76.0 | 78.5 | 78.5 |
|  | Other | 27.8 | 24.8 | 24.0 | 21.5 | 21.5 |
| Father's country of birth | Sweden | 70.7 | 74.7 | 74.7 | 77.3 | 77.6 |
|  | Other | 29.3 | 25.3 | 25.3 | 22.7 | 22.4 |
| Family disposable income quintile at birth | Lowest | 14.3 | 15.1 | 14.7 | 13.3 | 15.4 |
|  | Second | 21.7 | 20.4 | 20.8 | 19.4 | 19.1 |
|  | Third | 19.9 | 20.4 | 21.6 | 21.1 | 19.3 |
|  | Fourth | 22.3 | 22.3 | 21.6 | 22.6 | 22.1 |
|  | Highest | 21.8 | 21.8 | 21.4 | 23.6 | 24.2 |
| Parental educational attainment at birth | ≤9 years | 9.0 | 8.0 | 6.5 | 5.8 | 7.0 |
|  | 10 - 11 years | 42.0 | 43.3 | 40.5 | 39.9 | 40.7 |
|  | ≥13 years | 49.0 | 48.7 | 53.0 | 54.2 | 52.3 |
| Maternal age | ≤20 | 1.8 | 2.6 | 1.8 | 1.6 | 1.9 |
|  | 20-24 | 13.0 | 15.7 | 14.7 | 13.4 | 16.3 |
|  | 25-29 | 26.6 | 30.3 | 31.0 | 31.3 | 32.0 |
|  | 30-34 | 32.1 | 31.1 | 33.8 | 34.9 | 31.8 |
|  | 35-39 | 21.4 | 16.4 | 15.7 | 16.0 | 15.1 |
|  | 40+ | 5.2 | 4.0 | 3.2 | 2.8 | 2.9 |
| Paternal age | ≤20 | 0.7 | 0.8 | 0.5 | 0.5 | 0.4 |
|  | 20-24 | 7.2 | 8.9 | 7.2 | 7.0 | 8.2 |
|  | 25-29 | 20.8 | 24.2 | 23.5 | 23.5 | 25.0 |
|  | 30-34 | 30.8 | 31.5 | 33.5 | 33.7 | 32.5 |
|  | 35-39 | 22.8 | 20.4 | 22.0 | 22.0 | 21.4 |
|  | 40+ | 17.7 | 14.2 | 13.3 | 13.3 | 12.6 |

Table 1: continued
|  |  | Extremely<br>to very<br>preterm | Moderate<br>to late<br>preterm | Term | Post-term | Very<br>Post-term |
| --- | --- | --- | --- | --- | --- | --- |
| Gestational weeks |  | 21 - 31 | 32 - 36 | 37 - 41 | 42 | 43 - 45 |
| Number of observations |  | 2,601 | 20,271 | 438,215 | 34,828 | 3,706 |
| Percentage of the cohort |  | 0.5 | 4.1 | 87.7 | 7.0 | 0.7 |
|  |  | % | % | % | % | % |
| Maternal country of birth | Sweden | 72.2 | 75.2 | 76.0 | 78.5 | 78.5 |
|  | Other Nordic | 5.1 | 4.6 | 4.2 | 4.2 | 4.6 |
|  | Other European | 4.3 | 3.6 | 3.6 | 3.5 | 3.3 |
|  | Baltic States /Russia | 0.6 | 0.5 | 0.5 | 0.7 | 0.5 |
|  | Africa | 5.3 | 3.1 | 3.4 | 4.9 | 5.5 |
|  | Middle East | 7.3 | 6.7 | 7.0 | 4.8 | 4.5 |
|  | Asia /Oceania | 3.2 | 3.7 | 2.8 | 1.7 | 1.7 |
|  | North America | 0.5 | 0.6 | 0.6 | 0.6 | 0.4 |
|  | South America | 1.5 | 2.0 | 1.9 | 1.3 | 1.1 |
| Paternal country of birth | Sweden | 70.7 | 74.7 | 74.7 | 77.3 | 77.6 |
|  | Other Nordic | 4.3 | 3.5 | 3.2 | 3.0 | 3.4 |
|  | Other European | 4.6 | 4.5 | 4.7 | 4.6 | 4.5 |
|  | Baltic States /Russia | 0.2 | 0.2 | 0.2 | 0.2 | 0.2 |
|  | Africa | 6.2 | 4.1 | 4.1 | 5.7 | 5.8 |
|  | Middle East | 8.6 | 7.6 | 8.3 | 5.8 | 5.5 |
|  | Asia /Oceania | 2.4 | 2.3 | 1.9 | 1.2 | 1.1 |
|  | North America | 0.8 | 0.7 | 0.8 | 0.7 | 0.70 |
|  | South America | 2.3 | 2.3 | 2.2 | 1.5 | 1.21 |
| Intellectual disability |  | 5.6 | 1.8 | 0.9 | 1.0 | 1.6 |

Examining associations between gestational duration and risk of intellectual disability in a model using a continuous exposure variable with statistical adjustment for potential confounders (Figure 2), the adjusted odds ratio (aOR) for risk at extremely preterm birth (at 24 weeks) was estimated at 14.54 [95% CI 11.46 to 18.44]. This risk decreased with gestational age towards term (aOR_32 weeks_=3.59 [3.22 to 4.01]; aOR_37 weeks_=1.50 [1.38 to 1.63]; aOR_38 weeks_=1.26 [1.16 to 1.37]; aOR_39 weeks_=1.10 [1.04 to 1.17]) after which it increased with gestational age post-term (aOR_42 weeks_=1.16 [1.08 to 1.25]; aOR_43 weeks_=1.41 [1.21 to 1.64]; aOR_44 weeks_=1.71 [1.34 to 2.18]; aOR_45 weeks_=2.07 [1.47 to 2.92]).

**Fig 2:**
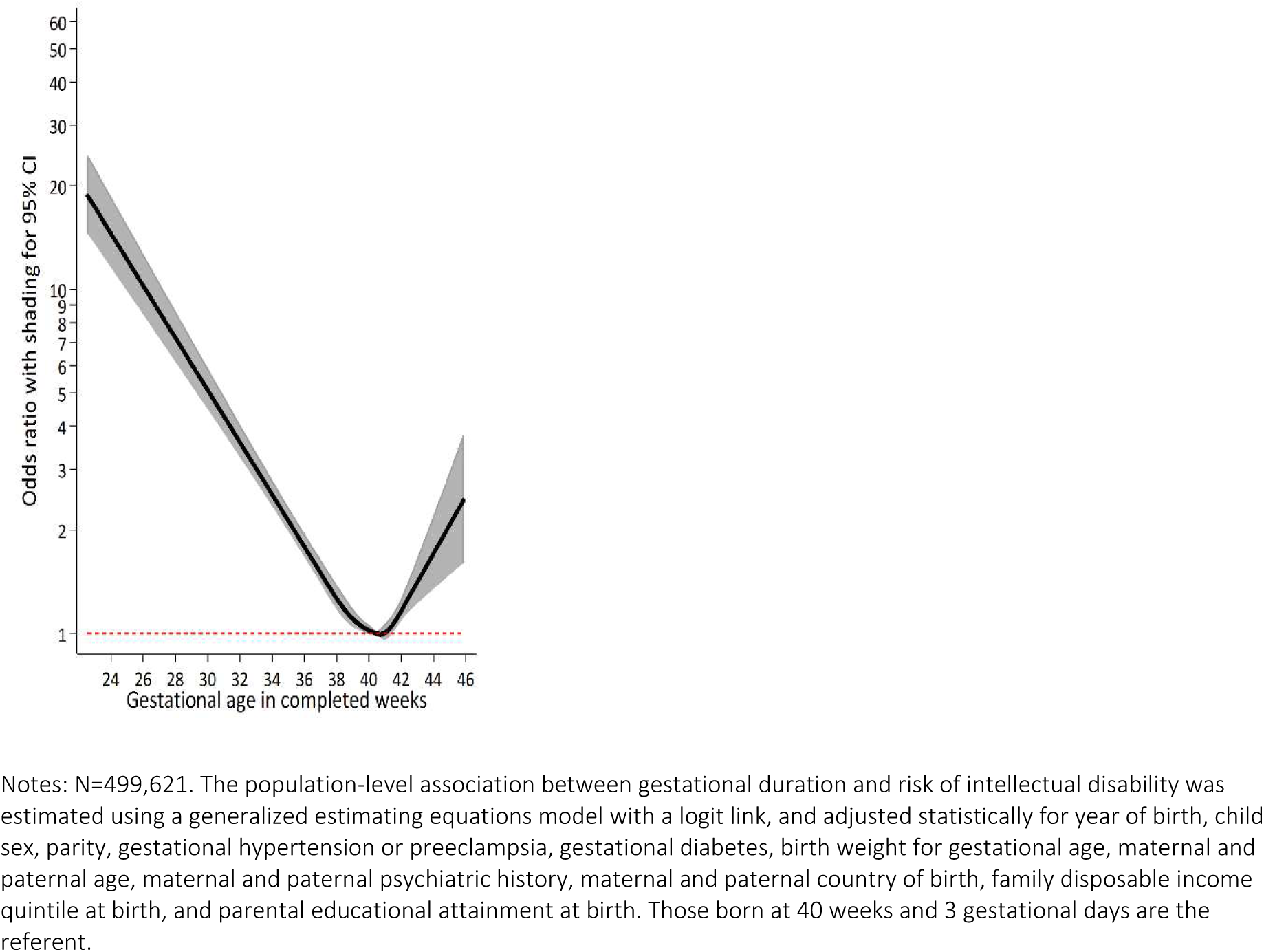
Population-level association between gestational duration and risk of intellectual disability.

We report associations using a categorical exposure variable in an online supplement (Table S7). Irrespective of gestational length, risk of intellectual disability was greatest among those showing evidence of fetal growth restriction (Table 2). This difference was most pronounced in the preterm group, but our results suggest risk of intellectual disability was also increased among children born post-term and growth-restricted. Associations between gestational length and risk of intellectual disability persisted when we repeated our analysis in a nested cohort of outcome-discordant siblings (Figure 3, Table S7).

**Fig 3:**
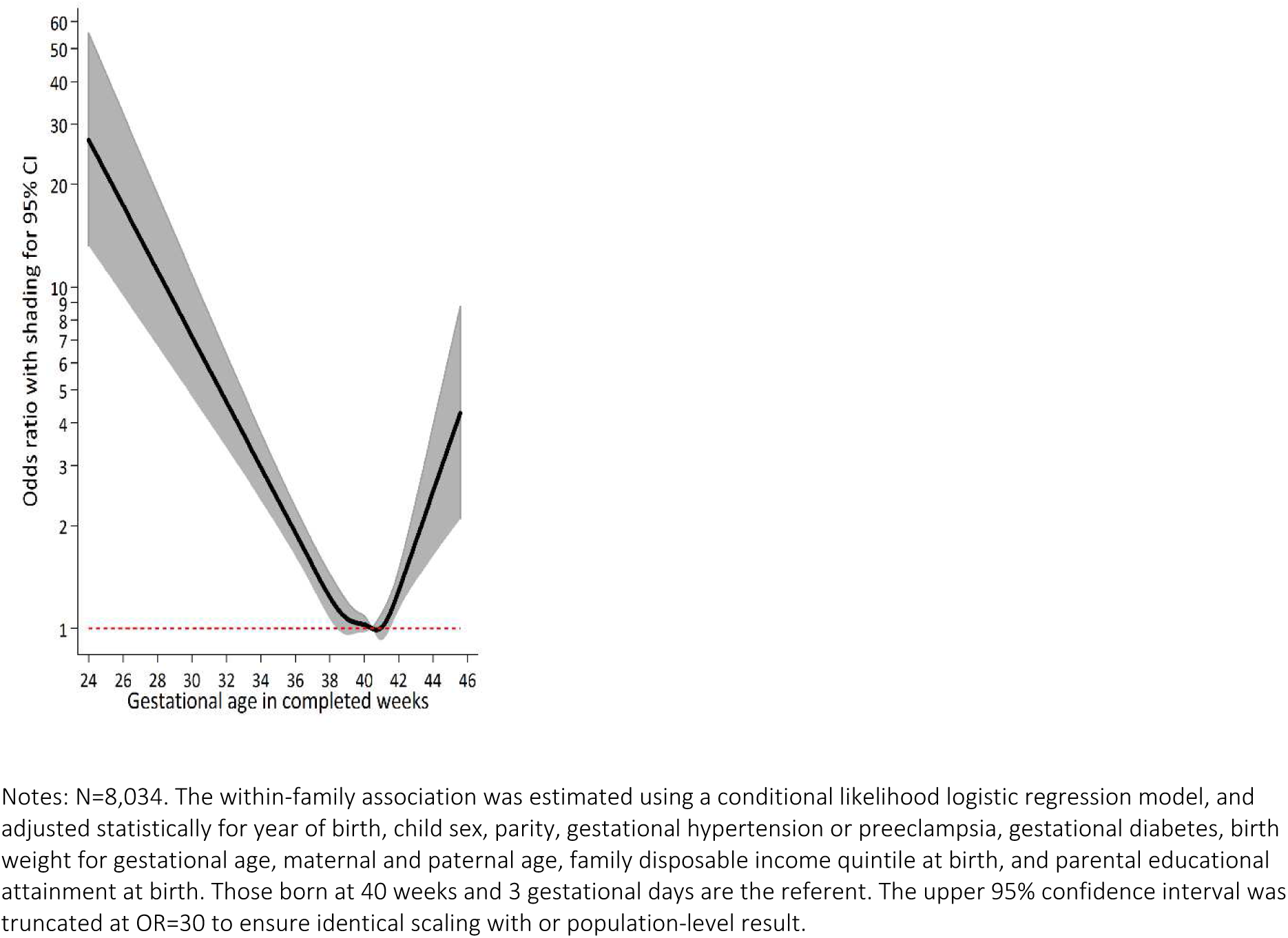
Within-family association between gestational duration and risk of intellectual disability.

**Table 2:** Interaction between gestational duration and fetal growth in relation to risk of intellectual disability.

| Gestational duration <sup>a</sup> /<br>weight-for-gestational age<br>category <sup>b</sup> | Odds<br>ratio <sup>c</sup> | 95% CI |  | *p* | n (%) <sup>d</sup> | N <sup>e</sup> | Percentage of<br>the cohort |
| --- | --- | --- | --- | --- | --- | --- | --- |
|  |  | (lower | upper) |  |  |  |  |
| Preterm / small | 3.77 | (2.97 | 4.84) | <0.001 | 73 (3.8) | 1,935 | 0.4 % |
| Preterm / appropriate | 2.24 | (2.01 | 2.51) | <0.001 | 380 (2.1) | 18,256 | 3.7 % |
| Preterm / large | 2.36 | (1.81 | 3.07) | <0.001 | 61 (2.3) | 2,681 | 0.5 % |
| Term / small | 1.88 | (1.73 | 2.05) | <0.001 | 731 (1.8) | 41,579 | 8.3 % |
| Term / appropriate | 1.00 |  |  |  | 3,008 (0.9) | 352,016 | 70.5 % |
| Term / large | 1.06 | (0.95 | 1.18) | 0.27 | 402 (0.9) | 44,620 | 8.9 % |
| Post-term / small | 2.29 | (1.83 | 2.85) | <0.001 | 85 (2.2) | 3,822 | 0.8 % |
| Post-term / appropriate | 1.11 | (0.98 | 1.25) | 0.10 | 292 (1.0) | 30,840 | 6.2 % |
| Post-term / large | 1.22 | (0.87 | 1.69) | 0.24 | 37 (1.0) | 3,872 | 0.8 % |

Notes: (a) Preterm was defined as birth < 37 completed weeks of gestation. Term birth was defined as birth between 37-41 completed weeks of gestation. Post-term birth was defined as birth at ≥42 completed weeks of gestation. (b) Fetal growth categories were defined as small-for-gestational age [in the lowest decile of the gestational age-specific birthweight distribution], appropriate-for-gestational age [in the 11<sup>th</sup> to 90<sup>th</sup> decile of the gestational age-specific birthweight distribution] and large-for-gestational age [in the upper decile of the gestational age-specific birthweight distribution]. (c) Population-level associations were estimated using a generalized estimating equations model with a logit link, and adjusted statistically for year of birth, child sex, parity, gestational hypertension or preeclampsia, gestational diabetes, maternal and paternal age, maternal and paternal psychiatric history, maternal and paternal country of birth, family disposable income quintile at birth, and parental educational attainment at birth. (d) Number and percentage of ID cases within gestational duration / fetal growth category; (e) Number of observations within gestational duration / fetal growth category. (e) N=499,621.

In a subset of the cohort without a diagnosis of ASD or ADHD, pre- and post-term birth remained associated with increased risk of intellectual disability (Figure S1, Table S3). Among those born at 21 to 31 completed weeks of gestation, risk of intellectual disability was lesser when the baby was delivered by Caesarean section, while Caesarean birth was associated with greater risk than vaginal birth between 37 to 41 weeks gestation (Table S4). There was no consistent variation in risk due to unassisted versus assisted delivery within gestational age categories (Table S5). Importantly, risk of intellectual disability associated with early or late birth remained when considering those born in vaginal or unassisted deliveries (Tables S4 and S5). Among those born between 37 and 41 weeks, risk of intellectual disability was greater when birth was induced (Table S6). This effect existed independently of the influence of fetal growth restriction or other potential confounders. Finally, children born in induced post-term deliveries were at greater risk of intellectual disability than children born spontaneously at term, while the increase in risk associated with spontaneous post-term birth was lesser (Table S6).

## Discussion

In this large population-based study, we found a greater risk of intellectual disability without a common genetic cause among preterm and post-term births compared with term births. These associations were evident in analyses using the full sample, as well as in a nested cohort of matched outcome-discordant siblings. Risk of intellectual disability was greatest among those showing evidence of fetal growth restriction, especially when born before or after term. To our knowledge, this is the first total-population study to estimate risk of intellectual disability without a common genetic cause over the entire range of gestation using high-quality prospectively measured data. In addition to a range of measured confounders, this study explored the influence of unmeasured familial effects using a matched sibling design. This allowed us to take into account unmeasured familial confounding of the association between intellectual disability and gestational length, as these traits are heritable within families^[10, 38, 39]^.

There were several limitations. First, 5 percent of the study cohort had missing data on gestational age at birth or other covariates. Although we cannot know with certainty how these exclusions may have affected our result, sensitivity analyses suggest that our estimates may have been conservative as they may have excluded preterm children with higher prevalence of intellectual disability (Table S2).

Second, we did not have information on whether gestational length was calculated by the mother’s report of her last menstrual period or based on ultrasound measurement in specific pregnancies. As our sample includes births from 1984 onwards, it is likely that there is greater measurement error in earlier cohort years, where gestational length would have been estimated on the basis of last menstrual period for a larger proportion of pregnancies. This may have resulted in overestimation of rates of post-term birth^[40-42]^ and underestimation of population-level^[43]^ and within-family associations^[44]^ between gestational length and later risk of intellectual disability. Third, while the matched-sibling design provides a powerful method to examine the influence of shared confounding, it is more sensitive than traditional methods to confounders not perfectly shared by the siblings.

Selection based on exposure-discordance could also prompt discordance in terms of non-shared confounding characteristics, which may bias the within-family effect^[44]^. The size and direction of such bias depends on the similarity or dissimilarity of matched siblings in terms of exposure and confounding characteristics^[44]^. Given that measurement error in the gestational age variable would have downwardly biased our estimate of the within-family effect, additional bias due to sibling non-shared confounding would have either offset this downward bias or further enhanced it. Fourth, there may be bias due to omitted non-shared confounding characteristics in our matched sibling analyses. For example, it is possible that prenatal infection^[45]^, maternal obesity^[46, 47]^, or use of drugs or alcohol^[48]^ may have influenced gestational length and resulted in greater risk of offspring intellectual disability in as far as these factors were present in one pregnancy but not the other.

The mechanisms underlying our findings are likely to differ depending on whether birth occurred before or after the due date. With regards to preterm birth, perturbations in development of the fetal brain because of shortened gestation can increase risk for longer-term neurodevelopmental problems^[49], [50]^. Our findings for preterm small-for-gestational age children would suggest that these effects might become particularly apparent if the fetus is already growth-restricted. After birth, further injury to the brain could result from respiratory support for preterm infants with immature pulmonary function ^[51]^. Mechanisms linking post-term birth with later risk of intellectual disability might involve placental deterioration or insufficiency causing fetal hypoxia or nutritional deficiencies ^[52]^, which in turn could result in injury to the fetal brain. Meconium aspiration, which is more common in post-term birth^[52]^, may result in neonatal asphyxia thereby incurring risk for brain injury and later neurodevelopmental problems^[53]^.

Our finding of associations among those born in unassisted or vaginal deliveries suggested that adverse obstetric circumstances did not explain the higher risk of intellectual disability associated with birth at <37 or >42 weeks. Furthermore, our findings suggest that risk of intellectual disability increases with induction of labor at further post-term gestation, although these estimates are likely to be biased by the higher risk nature of induced pregnancies as a whole (Table S6). Risk of intellectual disability may have also increased with advancing post-term gestational age when delivery started spontaneously, although our data may have been underpowered to detect these more subtle effects (Table S6). Importantly, given that the decision to induce labor will be informed by other factors than gestational length alone, we cannot infer from our data whether the risks associated with post-term delivery could be curtailed by induction of labor around term. This question may therefore be better answered by future research studies designed specifically to address this issue. Finally, in terms of the generalizability of our findings, the risks identified in our study may vary with regional differences in practice regarding the management of pre- or post-term pregnancy and in the quality of obstetric and neonatal care.

Our findings are consistent with other studies examining risk of cognitive deficit in relation to birth before or after term gestational duration ^[12-16, 54-60]^. These studies suggest there may be increased risk of intellectual disability^[12, 54]^, special educational needs^[14, 58, 59]^, poorer performance in school^[12, 15, 16, 55, 59]^ and lower IQ in childhood^[13, 60]^ or adulthood^[56, 57]^. The independent risk of intellectual disability associated with being born small for gestational age is consistent with earlier studies examining other outcomes for fetal growth-restriction in infants born at preterm or post-term gestational duration^[13, 15, 61, 62]^.

In conclusion, our findings suggest that delivery at non-optimal gestational age is associated with greater risk of intellectual disability in offspring in the absence of common genetic causes. This association existed independently of a range of measured potential confounders as well as unmeasured confounding from shared familial factors. While this study cannot provide conclusive evidence for causality, our use of a matched sibling design offered a stronger approach to dealing with confounding due to unmeasured shared familial factors, therefore providing a better estimate of the causal effect than studies using traditional methods for dealing with confounding. As birth at non-optimal gestational duration may be linked causally with greater risk of intellectual disability, it is important that the mechanisms underlying these associations are elucidated because of their relevance to clinical practice concerning elective delivery within the term period and the mitigation of risk in children who are born post-term.

---

**Table S1.** Excluded genetic and inborn metabolic syndromes.

|  | ICD-10<br>diagnostic code and description |  | ICD-9<br>diagnostic code and description |  | ICD-8<br>diagnostic code and description |  |
| --- | --- | --- | --- | --- | --- | --- |
| **Genetic defects** | Q85.0 | Neurofibromatosis (non-malignant) | 237.7 | Neurofibromatosis (uncertain behavior) | 743.4 | Neurofibromatosis |
|  | Q85.1 | Tuberous sclerosis | 759.5 | Tuberous sclerosis | 759.6 | Tuberous sclerosis |
|  | All Q90-Q99 | Chromosomal abnormalities, not elsewhere specified | 758 | Chromosomal anomalies | 759.3 | Down's syndrome |
|  |  |  |  |  | 759.4 | Other syndromes due to autosomal abnormality |
|  |  |  |  |  | 759.5 | Syndromes due to sex chromosome abnormality |
|  |  |  |  |  | 759.8 | Other specified syndromes |
|  |  |  |  |  | 759.9 | Multiple congenital anomalies, unspecified |
| **Inborn errors of metabolism** | All of E70-E72 | Metabolic disorders | All of 270 | Disorders of amino-acid transport and metabolism | All of 270 | Congenital disorders of amino-acid metabolism |

**Table S2:**
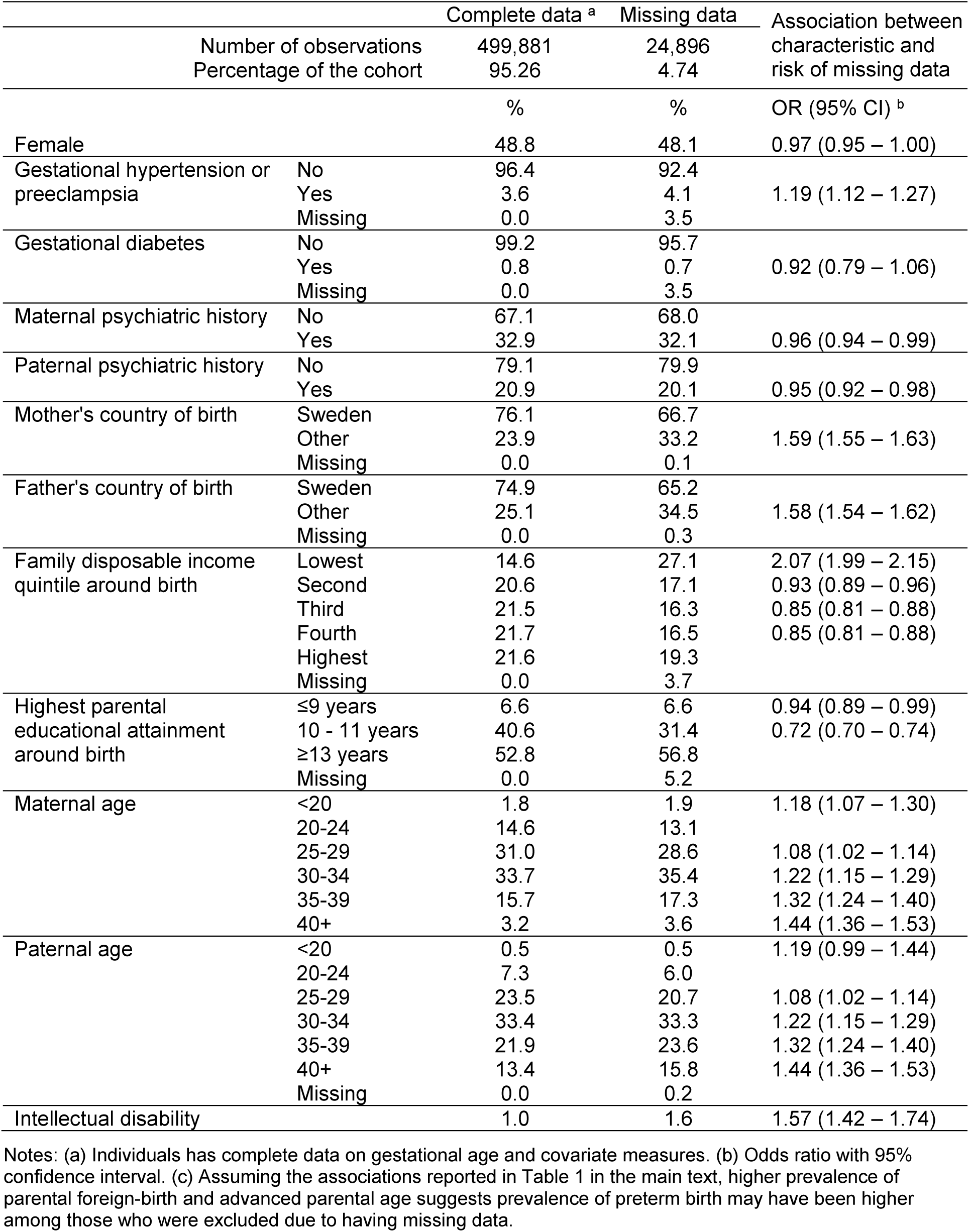
Characteristics of individuals with complete and missing data.

**Figure S1.**
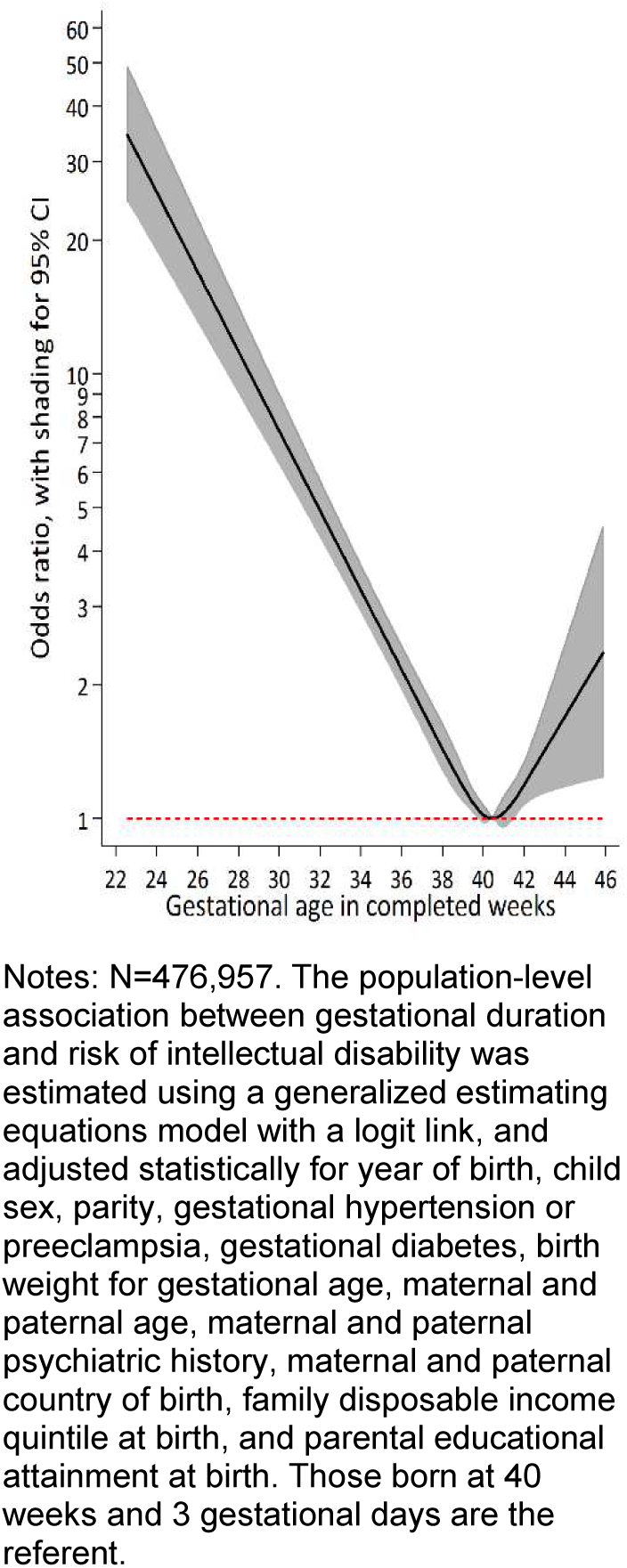
Population-level associations between risk of intellectual disability and gestational duration among those without ASD or ADHD.

**Table S3.** Population-level associations between risk of intellectual disability and gestational duration among those without ASD or ADHD.

| Number of completed weeks | Odds ratio | 95% CI (lower upper) | *p* | *n*<sup>3</sup> | *N*<sup>4</sup> |
| --- | --- | --- | --- | --- | --- |
| 21 - 31 | 8.25 | (6.49 10.48) | <0.001 | 79 | 2,323 |
| 32 - 36 | 2.22 | (1.90 2.60) | <0.001 | 185 | 19,073 |
| 37 - 41 | 1.00 |  |  | 1,694 | 418,827 |
| 42 | 1.07 | (0.90 1.27) | 0.44 | 139 | 33,227 |
| 43 - 45 | 1.68 | (1.13 2.49) | 0.011 | 25 | 3,507 |

Notes: N=476,957. The population-level association between gestational duration and risk of intellectual disability was estimated using a generalized estimating equations model with a logit link, and adjusted statistically for year of birth, child sex, parity, gestational hypertension or preeclampsia, gestational diabetes, birth weight for gestational age, maternal and paternal age, maternal and paternal psychiatric history, maternal and paternal country of birth, family disposable income quintile at birth, and parental educational attainment at birth. Those born between 37 and 41 completed weeks are the referent.

**Table S4.** Risk of intellectual disability among those born at varying gestational duration in vaginal or Caesarean deliveries.

|  |  | Odds ratio | 95% CI (lower upper) | *p* | *n*<sup>3</sup> | *N*<sup>4</sup> |
| --- | --- | --- | --- | --- | --- | --- |
| 21 - 31 weeks | Vaginal | 8.02 | (6.29 10.23) | <0.001 | 77 | 1,070 |
|  | Caesarean | 4.65 | (3.63 5.96) | <0.001 | 69 | 1,530 |
| 32 - 36 weeks | Vaginal | 1.68 | (1.46 1.93) | <0.001 | 221 | 13,699 |
|  | Caesarean | 2.17 | (1.82 2.58) | <0.001 | 147 | 6,572 |
| 37 - 41 weeks | Vaginal | 1.00 |  |  | 3,520 | 380,114 |
|  | Caesarean | 1.23 | (1.12 1.35) | <0.001 | 621 | 58,101 |
| 42 weeks | Vaginal | 1.09 | (0.96 1.22) | 0.18 | 294 | 29,077 |
|  | Caesarean | 1.36 | (1.06 1.76) | 0.017 | 62 | 5,751 |
| 43 - 45 weeks | Vaginal | 1.52 | (1.13 2.06) | 0.006 | 44 | 2,873 |
|  | Caesarean | 1.93 | (1.13 3.31) | 0.017 | 14 | 834 |

Notes: N=499,621. The population-level association between gestational duration and risk of intellectual disability was estimated using a generalized estimating equations model with a logit link, and adjusted statistically for year of birth, child sex, parity, gestational hypertension or preeclampsia, gestational diabetes, birth weight for gestational age, maternal and paternal age, maternal and paternal psychiatric history, maternal and paternal country of birth, family disposable income quintile at birth, and parental educational attainment at birth. Those born vaginally between 37 and 41 completed weeks are the referent.

**Table S5.** Risk of intellectual disability among those born at varying gestational duration in unassisted or assisted deliveries.

|  |  | Odds ratio | 95% CI<br>(lower upper) |  | p | n<sup>3</sup> | N<sup>4</sup> |
| --- | --- | --- | --- | --- | --- | --- | --- |
| 21 - 31 weeks | Unassisted | 5.81 | (4.86 | 6.93) | <0.001 | 144 | 2,564 |
|  | Forceps or ventouse | 6.89 | (1.66 | 28.51) | 0.008 | 2 | 36 |
| 32 - 36 weeks | Unassisted | 1.80 | (1.61 | 2.02) | <0.001 | 354 | 19,321 |
|  | Forceps or ventouse | 1.80 | (1.04 | 3.10) | 0.036 | 14 | 950 |
| 37 - 41 weeks | Unassisted | 1.00 |  |  |  | 3,814 | 403,120 |
|  | Forceps or ventouse | 1.12 | (0.99 | 1.25) | 0.065 | 327 | 35,095 |
| 42 weeks | Unassisted | 1.13 | (1.00 | 1.26) | 0.045 | 316 | 29,928 |
|  | Forceps or ventouse | 0.96 | (0.70 | 1.33) | 0.82 | 40 | 4,900 |
| 43 - 45 weeks | Unassisted | 1.51 | (1.13 | 2.01) | 0.006 | 48 | 3,154 |
|  | Forceps or ventouse | 2.03 | (1.08 | 3.81) | 0.027 | 10 | 553 |

Notes: N=499,621. The population-level association between gestational duration and risk of intellectual disability was estimated using a generalized estimating equations model with a logit link, and adjusted statistically for year of birth, child sex, parity, gestational hypertension or preeclampsia, gestational diabetes, birth weight for gestational age, maternal and paternal age, maternal and paternal psychiatric history, maternal and paternal country of birth, family disposable income quintile at birth, and parental educational attainment at birth. Those born between 37 and 41 completed weeks in unassisted deliveries are the referent.

**Table S6.** Risk of intellectual disability among those born at varying gestational duration in spontaneous or induced deliveries.

|  |  | Odds ratio | 95% CI<br>(lower upper) |  | p | n<sup>3</sup> | N<sup>4</sup> |
| --- | --- | --- | --- | --- | --- | --- | --- |
| 21 - 31 weeks | Spontaneous | 5.73 | (4.65 | 7.07) | <0.001 | 103 | 1,990 |
|  | Induced | 5.07 | (0.77 | 33.43) | 0.092 | 1 | 21 |
| 32 - 36 weeks | Spontaneous | 1.83 | (1.59 | 2.11) | <0.001 | 225 | 13,108 |
|  | Induced | 1.94 | (1.30 | 2.91) | 0.001 | 26 | 1,456 |
| 37 - 41 weeks | Spontaneous | 1.00 |  |  |  | 2,671 | 303,315 |
|  | Induced | 1.26 | (1.10 | 1.45) | <0.001 | 259 | 23,845 |
| 42 weeks | Spontaneous | 1.04 | (0.88 | 1.21) | 0.67 | 163 | 17,032 |
|  | Induced | 1.36 | (1.10 | 1.68) | 0.005 | 90 | 8,593 |
| 43 - 45 weeks | Spontaneous | 1.25 | (0.76 | 2.06) | 0.38 | 16 | 1,204 |
|  | Induced | 1.74 | (1.10 | 2.75) | 0.018 | 19 | 1,241 |

Notes: N=371,805. The population-level association between gestational duration and risk of intellectual disability was estimated using a generalized estimating equations model with a logit link, and adjusted statistically for year of birth, child sex, parity, gestational hypertension or preeclampsia, gestational diabetes, birth weight for gestational age, maternal and paternal age, maternal and paternal psychiatric history, maternal and paternal country of birth, family disposable income quintile at birth, and parental educational attainment at birth. Those born between 37 and 41 completed weeks in spontaneous deliveries are the referent.

**Table S7.** Population-level and within-family associations between gestational duration and risk of intellectual disability.

| Number of completed weeks | Population-level association <sup>a</sup> |  |  |  |  |  | Within-family association <sup>b</sup> |  |  |  |  |  |
| --- | --- | --- | --- | --- | --- | --- | --- | --- | --- | --- | --- | --- |
|  | Odds ratio | 95% CI<br>(lower upper) |  | *p* | *n* <sup>c</sup> | *N* <sup>d</sup> | Odds ratio | 95% CI<br>(lower upper) |  | *p* | *n* <sup>c</sup> | *N* <sup>d</sup> |
| 21 - 31 | 5.72 | (4.80 | 6.82) | <0.001 | 146 | 2,601 | 7.84 | (4.55 | 13.50) | <0.001 | 85 | 112 |
| 32 - 36 | 1.78 | (1.60 | 2.00) | <0.001 | 368 | 20,271 | 1.79 | (1.42 | 2.24) | <0.001 | 213 | 430 |
| 37 - 41 | 1.00 |  |  |  | 4,141 | 438,215 | 1.00 |  |  |  | 2,706 | 6,836 |
| 42 | 1.08 | (0.97 | 1.21) | 0.15 | 356 | 34,828 | 1.21 | (0.99 | 1.48) | 0.056 | 252 | 579 |
| 43 - 45 | 1.54 | (1.19 | 2.01) | 0.001 | 58 | 3,706 | 2.07 | (1.28 | 3.36) | 0.003 | 40 | 77 |

Notes: (a) Population-level associations were estimated using a generalized estimating equations model with a logit link, and adjusted statistically for year of birth, child sex, parity, gestational hypertension or preeclampsia, gestational diabetes, birth weight for gestational age, maternal and paternal age, maternal and paternal psychiatric history, maternal and paternal country of birth, family disposable income quintile at birth, and parental educational attainment at birth. (b) The within-family association was estimated using a conditional likelihood logistic regression model, and adjusted statistically for year of birth, child sex, parity, gestational hypertension or preeclampsia, gestational diabetes, birth weight for gestational age, maternal and paternal age, family disposable income quintile at birth, and parental educational attainment at birth. (c) Number of ID cases within gestational age category; (d) Number of observations within gestational age category.

## References

1. American Psychiatric Association. Diagnostic and Statistical Manual of Mental Disorders: DSM-IV. Washington DC.1994.

2. World Health Organisation. Definition: intellectual disability: WHO; 2016 [cited 2016 05/05/2016]. Available from: http://www.euro.who.int/en/health-topics/noncommunicable-diseases/mental-health/news/news/2010/15/childrens-right-to-family-life/definition-intellectual-disability

3. King BH, Toth KE, Hodapp RM, Dykens EM. Intellectual disability. In: Sadock BJ SV, Ruiz P, editor. Comprehensive textbook of psychiatry. 9 ed. Philadelphia: Lippincott Williams & Wilkins; 2009. p. 3444 – 74.

4. Lakhan R, Ekundayo OT, Shahbazi M. An estimation of the prevalence of intellectual disabilities and its association with age in rural and urban populations in India. J Neurosci Rural Pract. 2015;6(4):523–8. doi:10.4103/0976-3147.165392.

5. Maulik PK, Mascarenhas MN, Mathers CD, Dua T, Saxena S. Prevalence of intellectual disability: a meta-analysis of population-based studies. Res Dev Disabil. 2011;32(2):419–36. Epub 2011/01/18. doi:10.1016/j.ridd.2010.12.018. PubMed PMID: 21236634.

6. McKenzie K, Milton M, Smith G, Ouellette-Kuntz H. Systematic Review of the Prevalence and Incidence of Intellectual Disabilities: Current Trends and Issues. Curr Dev Disord Rep. 2016;3(2):104–15. doi:10.1007/s40474-016-0085-7.

7. Doran CM, Einfeld SL, Madden RH, Otim M, Horstead SK, Ellis LA, et al. How much does intellectual disability really cost? First estimates for Australia. J Intellect Dev Disabil. 2012;37(1):42–9. Epub 2012/02/22. doi:10.3109/13668250.2011.648609. PubMed PMID: 22339044.

8. Ali A, Hassiotis A, Strydom A, King M. Self stigma in people with intellectual disabilities and courtesy stigma in family carers: a systematic review. Res Dev Disabil. 2012;33(6):2122–40. Epub 2012/07/13. doi:10.1016/j.ridd.2012.06.013. PubMed PMID: 22784823.

9. Heslop P, Blair P, Fleming P, Hoghton M, Marriott A, Rull L. Confidential inquiry into premature deaths of people with learning disabilities (CIPOLD). Bristol: Norah Fry Centre for Disability Studies, 2013.

10. Shapiro BK, Batshaw ML. Intellectual disability. In: Kliegman RM, Behrman RE, Jenson HB, Stanton BF, editors. Nelson textbook of pediatrics. Philadelphia: Elsevier Saunders; 2011.

11. Huang J, Zhu T, Qu Y, Mu D. Prenatal, Perinatal and Neonatal Risk Factors for Intellectual Disability: A Systemic Review and Meta-Analysis. PLoS One. 2016;11(4):e0153655. Epub 2016/04/26. doi:10.1371/journal.pone.0153655. PubMed PMID: 27110944; PubMed Central PMCID: PMCPMC4844149.

12. Moster D, Lie RT, Markestad T. Long-term medical and social consequences of preterm birth. N Engl J Med. 2008;359(3):262–73. doi:10.1056/NEJMoa0706475. PubMed PMID: 18635431.

13. Yang S, Platt RW, Kramer MS. Variation in child cognitive ability by week of gestation among healthy term births. Am J Epidemiol. 2010;171(4):399–406. Epub 2010/01/19. doi:10.1093/aje/kwp413. PubMed PMID: 20080810; PubMed Central PMCID: PMCPMC3435092.

14. MacKay DF, Smith GC, Dobbie R, Pell JP. Gestational age at delivery and special educational need: retrospective cohort study of 407,503 schoolchildren. PLoS Med. 2010;7(6):e1000289. Epub 2010/06/15. doi:10.1371/journal.pmed.1000289. PubMed PMID: 20543995; PubMed Central PMCID: PMCPMC2882432.

15. Noble KG, Fifer WP, Rauh VA, Nomura Y, Andrews HF. Academic achievement varies with gestational age among children born at term. Pediatrics. 2012;130(2):e257–64. Epub 2012/07/04. doi:10.1542/peds.2011-2157. PubMed PMID: 22753563; PubMed Central PMCID: PMCPMC3408682.

16. Abel KM, Heuvelman H, Wicks S, Rai D, Emsley R, Gardner R, et al. Gestational age at birth and academic performance: population-based cohort study. Int J Epidemiol. 2016; Advance Access.

17. Terry AR, Barker FG, 2nd, Leffert L, Bateman BT, Souter I, Plotkin SR. Neurofibromatosis type 1 and pregnancy complications: a population-based study. Am J Obstet Gynecol. 2013;209(1):46.e1–8. Epub 2013/03/29. doi:10.1016/j.ajog.2013.03.029. PubMed PMID: 23535241.

18. Kase JS, Visintainer P. The relationship between congenital malformations and preterm birth. J Perinat Med. 2007;35(6):538–42. Epub 2007/12/07. doi:10.1515/jpm.2007.132. PubMed PMID: 18052839.

19. Idring S, Rai D, Dal H, Dalman C, Sturm H, Zander E, et al. Autism Spectrum Disorders in the Stockholm Youth Cohort: Design, Prevalence and Validity. PLoS One. 2012;7(7). doi:10.1371/journal.pone.0041280. PubMed PMID: 22911770; PubMed Central PMCID: PMCPMC3401114.

20. Zeitlin J, Saurel-Cubizolles M-J, de Mouzon J, Rivera L, Ancel P-Y, Blondel B, et al. Fetal sex and preterm birth: are males at greater risk? Hum Reprod. 2002;17(10):2762–8. doi:10.1093/humrep/17.10.2762.

21. Schempf AH, Branum AM, Lukacs SL, Schoendorf KC. Maternal age and parity-associated risks of preterm birth: differences by race/ethnicity. Paediatr Perinat Epidemiol. 2007;21(1):34–43. Epub 2007/01/24. doi:10.1111/j.1365-3016.2007.00785.x. PubMed PMID: 17239177.

22. Xiong X, Saunders LD, Wang FL, Demianczuk NN. Gestational diabetes mellitus: prevalence, risk factors, maternal and infant outcomes. International Journal of Gynecology & Obstetrics. 2001;75(3):221–8. doi:http://dx.doi.org/10.1016/S0020-7292(01)00496-9.

23. Ananth CV, Savitz DA, Luther ER, Bowes WA, Jr. Preeclampsia and preterm birth subtypes in Nova Scotia, 1986 to 1992. Am J Perinatol. 1997;14(1):17–23. Epub 1997/01/01. doi:10.1055/s-2007-994090. PubMed PMID: 9259891.

24. Li X, Sundquist J, Sundquist K. Immigrants and preterm births: a nationwide epidemiological study in Sweden. Matern Child Health J. 2013;17(6):1052–8. Epub 2012/07/27. doi:10.1007/s10995-012-1087-7. PubMed PMID: 22833337.

25. Morgan VA, Croft ML, Valuri GM, Zubrick SR, Bower C, McNeil TF, et al. Intellectual disability and other neuropsychiatric outcomes in high-risk children of mothers with schizophrenia, bipolar disorder and unipolar major depression. Br J Psychiatry. 2012;200(4):282–9. doi:10.1192/bjp.bp.111.093070.

26. Mannisto T, Mendola P, Kiely M, O’Loughlin J, Werder E, Chen Z, et al. Maternal psychiatric disorders and risk of preterm birth. Ann Epidemiol. 2016;26(1):14–20. Epub 2015/11/21. doi:10.1016/j.annepidem.2015.09.009. PubMed PMID: 26586549; PubMed Central PMCID: PMCPMC4688227.

27. Li X, Sundquist J, Kane K, Jin Q, Sundquist K. Parental occupation and preterm births: a nationwide epidemiological study in Sweden. Paediatr Perinat Epidemiol. 2010;24:555–63.

28. Zheng X, Chen R, Li N, Du W, Pei L, Zhang J, et al. Socioeconomic status and children with intellectual disability in China. J Intellect Disabil Res. 2011;56(2):212–20.

29. Ruiz M, Goldblatt P, Morrison J, Kukla L, Švancara J, Riitta-Järvelin M, et al. Mother’s education and the risk of preterm and small for gestational age birth: a DRIVERS meta-analysis of 12 European cohorts. J Epidemiol Community Health. 2015. doi:10.1136/jech-2014-205387.

30. Stata. xtgee - Fit population-averaged panel-data models by using GEE: Stata; [10/10/2016]. Available from: http://www.stata.com/manuals13/xtxtgee.pdf.

31. Stata. mkspline - Linear and restricted cubic spline construction: Stata; [10/10/2016]. Available from: http://www.stata.com/manuals13/rmkspline.pdf.

32. Lahey BB, D’Onofrio BM. All in the Family: Comparing Siblings to Test Causal Hypotheses Regarding Environmental Influences on Behavior. Curr Dir Psychol Sci. 2010;19(5):319–23. Epub 2010/10/01. doi:10.1177/0963721410383977. PubMed PMID: 23645975; PubMed Central PMCID: PMCPMC3643791.

33. Shea V, Mesibov G. Adolescents and adults with autism. In: Volkmar FR, Paul R, Klin A, Cohen D, editors. Handbook of autism and pervasive developmental disorders. 5 ed: John Wiley & Sons; 2005.

34. Chakrabarti S, Fombonne E. Pervasive developmental disorders in preschool children: confirmation of high prevalence. Am J Psychiatry. 2005;162(6):1133–41. doi:10.1176/appi.ajp.162.6.1133. PubMed PMID: 15930062.

35. Voigt RG, Barbaresi WJ, Colligan RC, Weaver AL, Katusic SK. Developmental dissociation, deviance, and delay: Occurrence of attention-deficit-hyperactivity disorder in individuals with and without borderline-to-mild intellectual disability. Dev Med Child Neurol. 2006;48(10):831–5. Epub 2006/09/19. doi:10.1017/s0012162206001782. PubMed PMID: 16978463.

36. Leavey A, Zwaigenbaum L, Heavner K, Burstyn I. Gestational age at birth and risk of autism spectrum disorders in Alberta, Canada. J Pediatr. 2013;162(2):361–8. Epub 2012/09/06. doi:10.1016/j.jpeds.2012.07.040. PubMed PMID: 22947654.

37. Sucksdorff M, Lehtonen L, Chudal R, Suominen A, Joelsson P, Gissler M, et al. Preterm Birth and Poor Fetal Growth as Risk Factors of Attention-Deficit/Hyperactivity Disorder. Pediatrics. 2015. doi:10.1542/peds.2015-1043.

38. Kaufman L, Ayub M, Vincent JB. The genetic basis of non-syndromic intellectual disability: a review. J Neurodev Disord. 2010;2(4):182–209. doi:10.1007/s11689-010-9055-2. PubMed PMID: PMC2974911.

39. Lie RT, Wilcox AJ, Skjaerven R. Maternal and paternal influences on length of pregnancy. Obstet Gynecol. 2006;107(4):880–5. Epub 2006/04/04. doi:10.1097/01.aog.0000206797.52832.36. PubMed PMID: 16582127.

40. Haglund B. Birthweight distributions by gestational age: comparison of LMP-based and ultrasound-based estimates of gestational age using data from the Swedish Birth Registry. Paediatr Perinat Epidemiol. 2007;21:72–8. doi:10.1111/j.1365-3016.2007.00863.x.

41. Dietz PM, England LJ, Callaghan WM, Pearl M, Wier ML, Kharrazi M. A comparison of LMP-based and ultrasound-based estimates of gestational age using linked California livebirth and prenatal screening records. Paediatr Perinat Epidemiol. 2007;21:62–71. doi:10.1111/j.1365-3016.2007.00862.x.

42. Hoffman CS, Messer LC, Mendola P, Savitz DA, Herring AH, Hartmann KE. Comparison of gestational age at birth based on last menstrual period and ultrasound during the first trimester. Paediatr Perinat Epidemiol. 2008;22(6):587–96. doi:10.1111/j.1365-3016.2008.00965.x.

43. D Thomas, D Stram a, Dwyer J. Exposure Measurement Error: Influence on Exposure-Disease Relationships and Methods of Correction. Annu Rev Public Health. 1993;14(1):69–93. doi: doi:10.1146/annurev.pu.14.050193.000441. PubMed PMID: 8323607.

44. Frisell T, Oberg S, Kuja-Halkola R, Sjolander A. Sibling comparison designs: bias from non-shared confounders and measurement error. Epidemiology. 2012;23(5):713–20. Epub 2012/07/12. doi:10.1097/EDE.0b013e31825fa230. PubMed PMID: 22781362.

45. Dammann O, Kuban KC, Leviton A. Perinatal infection, fetal inflammatory response, white matter damage, and cognitive limitations in children born preterm. Mental retardation and developmental disabilities research reviews. 2002;8(1):46–50. Epub 2002/03/29. doi:10.1002/mrdd.10005. PubMed PMID: 11921386.

46. Sharashova EE, Anda EE, Grjibovski AM. Early pregnancy body mass index and spontaneous preterm birth in Northwest Russia: a registry-based study. BMC Pregnancy Childbirth. 2014;14(1):303. doi:10.1186/1471-2393-14-303.

47. Basatemur E, Gardiner J, Williams C, Melhuish E, Barnes J, Sutcliffe A. Maternal Prepregnancy BMI and Child Cognition: A Longitudinal Cohort Study. Pediatrics. 2013;131(1):56–63. doi:10.1542/peds.2012-0788.

48. Forray A. Substance use during pregnancy. F1000Research. 2016;|p5. doi:10.12688/f1000research.7645.1. PubMed PMID: 27239283; PubMed Central PMCID: PMCPMC4870985.

49. Ortinau C, Neil J. The neuroanatomy of prematurity: normal brain development and the impact of preterm birth. Clin Anat. 2015;28(2):168–83. Epub 2014/07/22. doi:10.1002/ca.22430. PubMed PMID: 25043926.

50. Pavlova MA, Krageloh-Mann I. Limitations on the developing preterm brain: impact of periventricular white matter lesions on brain connectivity and cognition. Brain. 2013;136(Pt 4):998–1011. Epub 2013/04/04. doi:10.1093/brain/aws334. PubMed PMID: 23550112.

51. Polglase GR, Miller SL, Barton SK, Kluckow M, Gill AW, Hooper SB, et al. Respiratory support for premature neonates in the delivery room: effects on cardiovascular function and the development of brain injury. Pediatr Res. 2014;75(6):682–8. Epub 2014/03/13. doi:10.1038/pr.2014.40. PubMed PMID: 24614803.

52. Galal M, Symonds I, Murray H, Petraglia F, Smith R. Postterm pregnancy. Facts, views & vision in ObGyn. 2012;4(3):175–87. Epub 2012/01/01. PubMed PMID: 24753906; PubMed Central PMCID: PMCPMC3991404.

53. Gulmezoglu AM, Crowther CA, Middleton P, Heatley E. Induction of labour for improving birth outcomes for women at or beyond term. Cochrane Database Syst Rev. 2012;(6):Cd004945. Epub 2012/06/15. doi:10.1002/14651858.CD004945.pub3. PubMed PMID: 22696345; PubMed Central PMCID: PMCPMC4065650.

54. Larroque B, Ancel PY, Marret S, Marchand L, Andre M, Arnaud C, et al. Neurodevelopmental disabilities and special care of 5-year-old children born before 33 weeks of gestation (the EPIPAGE study): a longitudinal cohort study. Lancet. 2008;371(9615):813–20. Epub 2008/03/11. doi:10.1016/s0140-6736(08)60380-3. PubMed PMID: 18328928.

55. Lipkind HS, Slopen ME, Pfeiffer MR, McVeigh KH. School-age outcomes of late preterm infants in New York City. Am J Obstet Gynecol. 2012;206(3): 222.e1-6. Epub 2012/03/03. doi:10.1016/j.ajog.2012.01.007. PubMed PMID: 22381605.

56. Ekeus C, Lindstrom K, Lindblad F, Rasmussen F, Hjern A. Preterm birth, social disadvantage, and cognitive competence in Swedish 18- to 19-year-old men. Pediatrics. 2010;125(1):e67–73. Epub 2009/12/09. doi:10.1542/peds.2008-3329. PubMed PMID: 19969613.

57. Eide MG, Oyen N, Skjaerven R, Bjerkedal T. Associations of birth size, gestational age, and adult size with intellectual performance: evidence from a cohort of Norwegian men. Pediatr Res. 2007;62(5):636–42. Epub 2007/09/07. doi:10.1203/PDR.0b013e31815586e9. PubMed PMID: 17805203.

58. Morse SB, Zheng H, Tang Y, Roth J. Early school-age outcomes of late preterm infants. Pediatrics. 2009;123(4):e622–9. Epub 2009/04/02. doi:10.1542/peds.2008-1405. PubMed PMID: 19336353.

59. Chyi LJ, Lee HC, Hintz SR, Gould JB, Sutcliffe TL. School outcomes of late preterm infants: special needs and challenges for infants born at 32 to 36 weeks gestation. J Pediatr. 2008;153(1):25–31. Epub 2008/06/24. doi:10.1016/j.jpeds.2008.01.027. PubMed PMID: 18571530.

60. Talge NM, Holzman C, Wang J, Lucia V, Gardiner J, Breslau N. Late-preterm birth and its association with cognitive and socioemotional outcomes at 6 years of age. Pediatrics. 2010;126(6):1124–31. Epub 2010/11/26. doi:10.1542/peds.2010-1536. PubMed PMID: 21098151.

61. Morken NH, Klungsoyr K, Skjaerven R. Perinatal mortality by gestational week and size at birth in singleton pregnancies at and beyond term: a nationwide population-based cohort study. BMC Pregnancy Childbirth. 2014;14:172. Epub 2014/06/03. doi:10.1186/1471-2393-14-172. PubMed PMID: 24885576; PubMed Central PMCID: PMCPMC4037279.

62. Abel KM, Dalman C, Svensson AC, Susser E, Dal H, Idring S, et al. Deviance in fetal growth and risk of autism spectrum disorder. Am J Psychiatry. 2013;170(4):391–8. Epub 2013/04/03. doi:10.1176/appi.ajp.2012.12040543. PubMed PMID: 23545793.

